# Chemically-induced epimutagenesis allows bypassing reproductive barriers in hybrid seeds

**DOI:** 10.1101/2021.07.23.453551

**Authors:** Jonathan Huc, Katarzyna Dziasek, Kannan Pachamuthu, Tristan Woh, Claudia Köhler, Filipe Borges

## Abstract

The “triploid block” prevents interploidy hybridizations in flowering plants, and is characterized by failure in endosperm development, arrest in embryogenesis, and seed collapse. Many genetic components of triploid seed lethality have been successfully identified in the model plant *Arabidopsis thaliana*, most notably the paternally expressed imprinted genes (PEGs) that are up-regulated in the tetraploid endosperm with paternal excess. Previous studies have shown that the paternal epigenome is a key determinant of the triploid block response, as the loss of DNA methylation in diploid pollen suppresses the triploid block almost completely. Here, we demonstrate that triploid seed collapse is bypassed in Arabidopsis plants treated with the DNA methyltransferase inhibitor 5-Azacytidine during seed germination and early growth. We have identified strong suppressor lines showing stable transgenerational inheritance of hypomethylation in CG context, as well as normalized expression of PEGs in triploid seeds. Importantly, differentially methylated loci segregate in the progeny of “epimutagenized” plants, which may allow the identification of epialleles involved in the triploid block response in future studies. Finally, we demonstrate that chemically-induced epimutagenesis allows bypassing hybridization barriers in crosses between different Capsella species, thus potentially emerging as a novel strategy for producing triploids and interspecific hybrids with high agronomical interest.

**One sentence summary:** Genome-wide loss of DNA methylation induced by 5-Azacytidine allows bypassing interploidy and interspecific hybridization barriers in Arabidopsis and Capsella.

## Introduction

Early studies in plants showed some of the first evidence of distinctive phenotypes dependent on the nature and dosage of parental chromosomes (Belling and Blakeslee, 1923; Blakeslee et al., 1920). This phenomenon, currently known as heterosis, is often observed in interploid and interspecific hybrids that display phenotypic values exceeding those in their parents, but its genetic and epigenetic basis remain poorly understood (Birchler et al., 2010; Hochholdinger and Baldauf, 2018). Plant breeders have been exploiting heterosis for thousands of years to create elite varieties of domesticated crops with enhanced growth and yield. However, more progress has been hindered by the existence of strong reproductive barriers that prevent heterosis, and a faster introgression of valuable alleles from wild species into domesticated cultivars (Kaneko and Bang, 2014).

In many angiosperms, interploidy crosses between diploid females and tetraploid males lead to abnormal endosperm development and seed collapse, which is known as the “triploid block” response (Köhler et al., 2021). The endosperm of most flowering plants is an essential triploid tissue that nourishes early embryo development in the seed, and is formed in the embryo sac by fertilization of the diploid central cell by an haploid sperm cell (Dresselhaus et al., 2016). Endosperm cellularization is a critical event for seed development, and relies on the correct balance between the dosage of maternal and paternal genomes (2m:1p), and distinct epigenetic features that are pre-established in the gametes to control genomic imprinting after fertilization (Gehring and Satyaki, 2017; Kawashima and Berger, 2014). Maternally expressed genes (MEGs) encode components of the Polycomb Repressive Complex 2 (PRC2), which is essential to silence the maternal allele of paternally expressed genes (PEGs) via deposition of tri-methylation at lysine 27 of histone 3 (H3K27me3) (Gehring and Satyaki, 2017; Köhler and Lafon-Placette, 2015). For a long time, this mechanism was sufficient to explain how imprinted gene expression is established in the endosperm. However, additional models emerged with the discovery of a paternally-inherited small RNA-directed DNA methylation (RdDM) pathway that is also involved in genomic imprinting (Vu et al., 2013; Calarco et al., 2012).

The triploid block traces back to the Endosperm Balance Number (EBN) hypothesis, which was developed in the early 80’s in potato and then extended to many other crops (Johnston and Hanneman, 1982; Ehlenfeldt and Hanneman, 1984). The EBN, or “effective ploidy”, is the ratio between maternal and paternal chromosomes or genetic factors required for development of a normal seed (Carputo et al., 1999, 1997; Johnston and Hanneman, 1982; Ehlenfeldt and Hanneman, 1984). These studies are highly relevant for plant breeding, as the success of interspecific and intergeneric hybridizations may be predicted and manipulated based on the EBN of each parent (Tonosaki et al., 2018), although this system seems to be restricted to only certain genera (Carputo et al., 1999). Additional breeding strategies to overcome hybridization barriers require ovule/ovary cultures and embryo rescue techniques prior to seed collapse, which have limited efficiency depending on the species (Cisneros and Tel-Zur, 2010; Sauer and Friml, 2008; Eeckhaut et al., 2006). In the model plant *Arabidopsis thaliana,* loss-of-function mutations in PEGs are able to suppress the triploid block (Kradolfer et al., 2013; Batista et al., 2019; Wolff et al., 2015). This clearly indicates that endosperm failure during interploidy hybridization results from up-regulation of PEGs, which is also observed in crosses between different Arabidopsis species (Kirkbride et al., 2015; Josefsson et al., 2006), and may have role in establishing interspecific hybridization barriers as well (Lafon-Placette et al., 2018). However, its potential application in plant breeding is limited, as imprinted genes are generally not well conserved (Kradolfer et al., 2013; Rodrigues and Zilberman, 2015), in a way that studies in Arabidopsis cannot be directly tested in crops. More recently, several studies have shown that the paternal epigenome triggers the triploid block response in Arabidopsis, which provided new ideas for plant breeding. Loss of DNA and histone methylation, as well as small-interfering RNAs (siRNAs) in diploid pollen restored viability of triploid seeds almost completely (Martinez et al., 2018; Borges et al., 2018; Erdmann et al., 2017; Schatlowski et al., 2014; Jiang et al., 2017; Satyaki and Gehring, 2019), suggesting that the paternal epigenome mediates genomic imprinting and endosperm balance in the developing seed.

Here, we show that transient genome-wide epimutagenesis induced by the chemical inhibitor of DNA methylation 5-Azacytidine allows bypassing the triploid block in Arabidopsis. We identified and characterized strong epigenetic suppressors of the triploid block response, showing stable transgenerational loss of CG methylation, and down-regulation of imprinted genes that are well-known triggers of triploid seed collapse. Finally, we demonstrate that epimutagenesis induced by 5-Azacytidine allows bypassing hybridization barriers in crosses between Capsella species, thus potentially emerging as a new method to facilitate the production of triploid plants and interspecific hybrids with high socio-economical interest for agriculture and crop improvement.

## Results

### Exposure to 5-Azacytidine during seed germination and early growth allows bypassing the triploid block

Chemical inhibition of DNA methyltransferases has been widely used to study the function of DNA methylation in several plant systems (Pecinka and Liu, 2014). For instance, cytosine analogs such as 5-Azacytidine and Zebularine are incorporated into newly replicated DNA, but do not become methylated (Pecinka and Liu, 2014), in a way that DNA methylation is passively erased during cell divisions (Jones and Taylor, 1980; Creusot et al., 1982; Santi et al., 1984). Recent whole-genome analysis of DNA methylation in Arabidopsis has shown that both chemicals lead to widespread loss of DNA methylation in all sequence contexts, and in a dose-dependent manner (Griffin et al., 2016), which can be stably inherited into subsequent generations or fully restored before fertilization (Baubec et al., 2009; Akimoto et al., 2007).

In order to test if epigenetic variation induced by 5-Azacytidine allows suppressing the triploid block response, we used the Arabidopsis mutant *jason* (Storme and Geelen, 2011; Erilova et al., 2009). In our growth conditions, plants homozygous for the *jas-3* allele in Col-0 background produce 30-40% of diploid pollen (Supplemental Figure 1), while the female gametophyte is haploid. This way, the triploid block response may be quantified after self-fertilization. In untreated *jas-3* plants and DMSO-treated controls, triploid seed abortion varied between 30 and 40% (Figure 1A), thus reflecting the amount of diploid pollen in these plants (Supplemental Figure 1). Strikingly, plants treated with different concentrations of 5-Azacytidine showed a variable and dose-dependent effect in the triploid block response (Figure 1A), which was significantly reduced to ~20% in plants treated with 100μg/mL of the chemical (Figure 1A). In particular lines (e.g. Aza1), seed collapse was reduced to less than 10% (Figure 1A), while the amount of diploid pollen remained unchanged (Supplemental Figure 1), thus suggesting strong suppression of the triploid block response. Close inspection of the seed set showed the presence of enlarged seeds (Figure 1B) that were routinely confirmed to be triploids by flow cytometric analysis of ploidy (Supplemental Figure 2).

**Figure 1.**
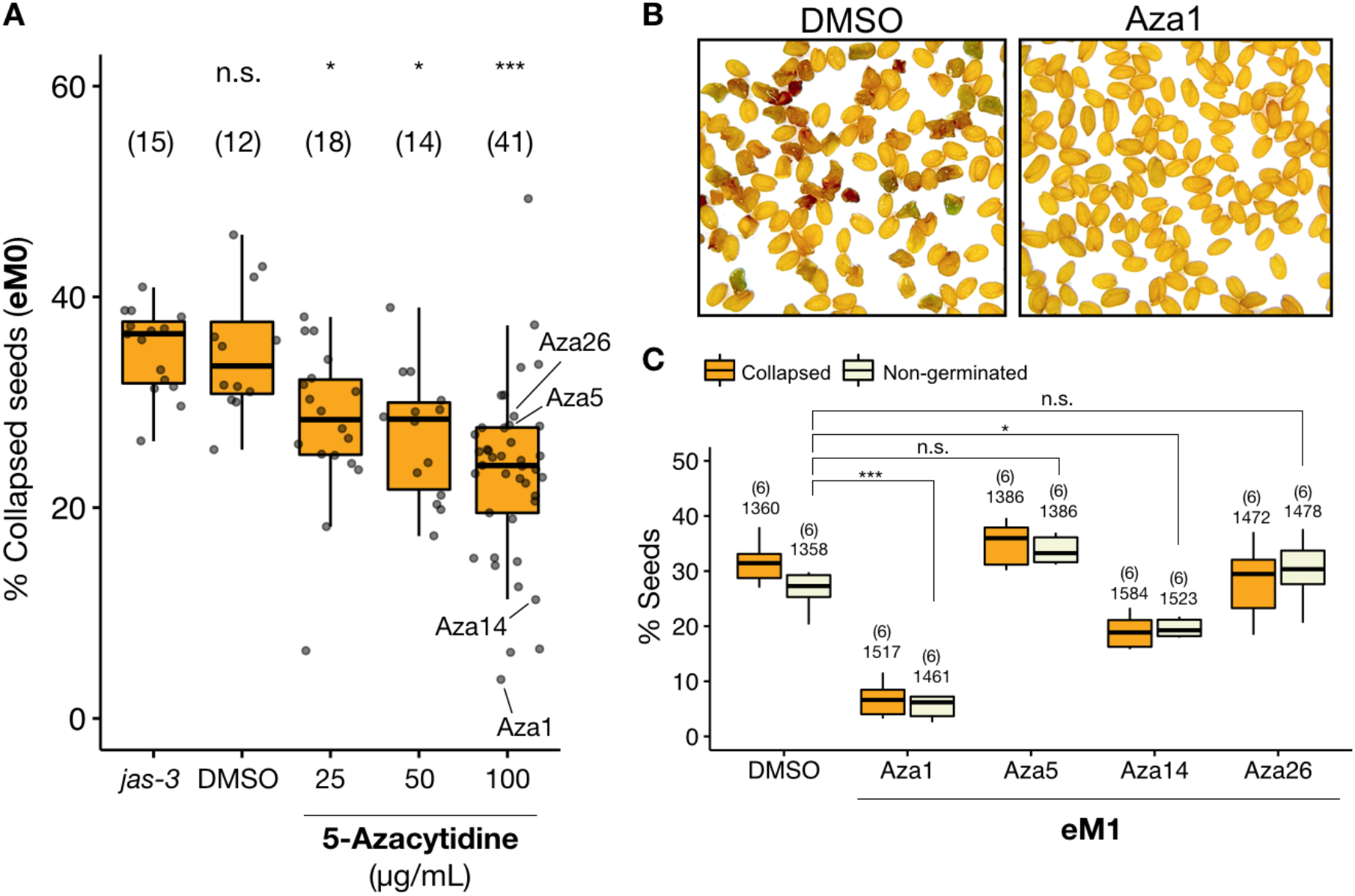
The triploid block is suppressed in *jas-3* plants treated with 5-Azacytidine. **(A)** The triploid block response was quantified by counting the number of aborted seeds in five siliques of untreated *jas-3* plants and DMSO controls, and plants exposed to three different concentrations of 5-Azacytidine (25, 50 and 100 μg/mL). Numbers on top of each box represent the number of individual plants used. Statistically significant difference in the percentage of collapsed seeds was calculated by ANOVA with a post hoc Dunnett test, using *jas-3* as the reference group (n.s. is not significant, * is *p* ≤ 0.05, and *** is *p* ≤ 0.001). **(B)** Representative images of seeds from DMSO controls and the strong suppressor Aza1, showing a decrease in the level of seed abortion in the suppressor line. **(C)** Seed abortion and germination assays were performed in the first generation after treatment with 5-Azacytidine (eM1) to evaluate the transgenerational stability of the suppressive effect. Numbers on top of each box indicate the number of siblings (top) and total number of seeds (bottom) counted. Statistically significant difference in the percentage of non-germinated seeds was calculated by ANOVA with a post hoc Dunnett test, using DMSO as the reference group (n.s. is not significant, * is *p* ≤ 0.05, and *** is *p* ≤ 0.001).

In Arabidopsis, known epigenetic suppressors of the triploid block show a suppressive effect only when paternally inherited (Schatlowski et al., 2014; Borges et al., 2018; Martinez et al., 2018; Satyaki and Gehring, 2019). We therefore asked if there is also a parental effect in triploid block suppression caused by 5-Azacytidine treatment, and tested this hypothesis by performing reciprocal crosses between DMSO control plants and the strong suppressor line Aza1. Indeed, we found that the suppressive effect caused by 5-Azacytidine is transmitted via the paternal genome (Supplemental Figure 3).

We then selected diploid seeds from individual 5-Azacytidine-treated lines to inspect the stability of the suppressive effect in the next generation (Supplemental Figure 4), which will be hereafter designated as eM1 (epimutagenized population 1). Seeds from six individual plants of two suppressor lines (Aza1 and Aza14) and two non-suppressors (Aza5 and Aza26) were compared to the DMSO controls, to show that seed abortion and germination rates were similar to those observed in eM0 plants that had been directly exposed to the chemical (Figure 1C). Taken together, these results demonstrate that exposing *jas-3* plants to 5-Azacytidine during seed germination and early growth allows bypassing the triploid block response at variable levels, in a dose-dependent and transgenerational manner.

### Genome-wide CG hypomethylation observed in suppressor lines

Whole-genome bisulfite sequencing (WGBS) was performed to test whether suppression of the triploid block in 5-Azacytidine-treated *jas-3* plants correlated with the loss of DNA methylation (Supplemental Data Set 1). Therefore, comparative methylome analyses were performed between the strong suppressor lines Aza1 (3,7% collapsed seeds), Aza14 (11,3%), Aza18 (6,6%) and Aza25 (6,3%), and the non-suppressor lines Aza5 (27,8%), Aza10 (30,7%), Aza16 (26,9%) and Aza26 (28,7%) that showed levels of seed collapse closer to the controls *jas-3* (mean 33,7%) and DMSO (mean 33,4%). Indeed, in suppressor lines, strong loss of CG methylation was observed at protein-coding genes and transposable elements (TEs), as compared to untreated *jas-3* and DMSO controls (Figure 2A and Supplemental Figure 5). Analysis of differentially methylated regions (DMRs) allowed the identification of 6393 hypomethylated CG DMRs among all suppressor lines (Figure 2B), which occurred primarily at protein-coding genes (Figure 2C and Supplemental Data Set 2). Interestingly, this analysis also showed variable patterns of hypomethylation in the different suppressors (Figure 2B), as the overlap between the three strongest suppressors (Aza1, Aza18 and Aza25) is limited to ~1/10 of the amount of DMRs detected in each individual line (Figure 2D). In contrast, CG methylation levels were mostly unchanged in all non-suppressors (Figure 2A and Supplemental Figure 5), confirming that genome-wide levels of CG methylation correlate with the triploid block response. Importantly, cytosine methylation in CHG and CHH contexts was mostly unchanged in both suppressor and non-suppressor lines (Supplemental Figure 5 and Supplemental Data Set 1). This indicates that non-CG methylation was rapidly restored in the first generation after 5-Azacytidine treatment, and is likely not responsible for the suppressive effect.

**Figure 2.**
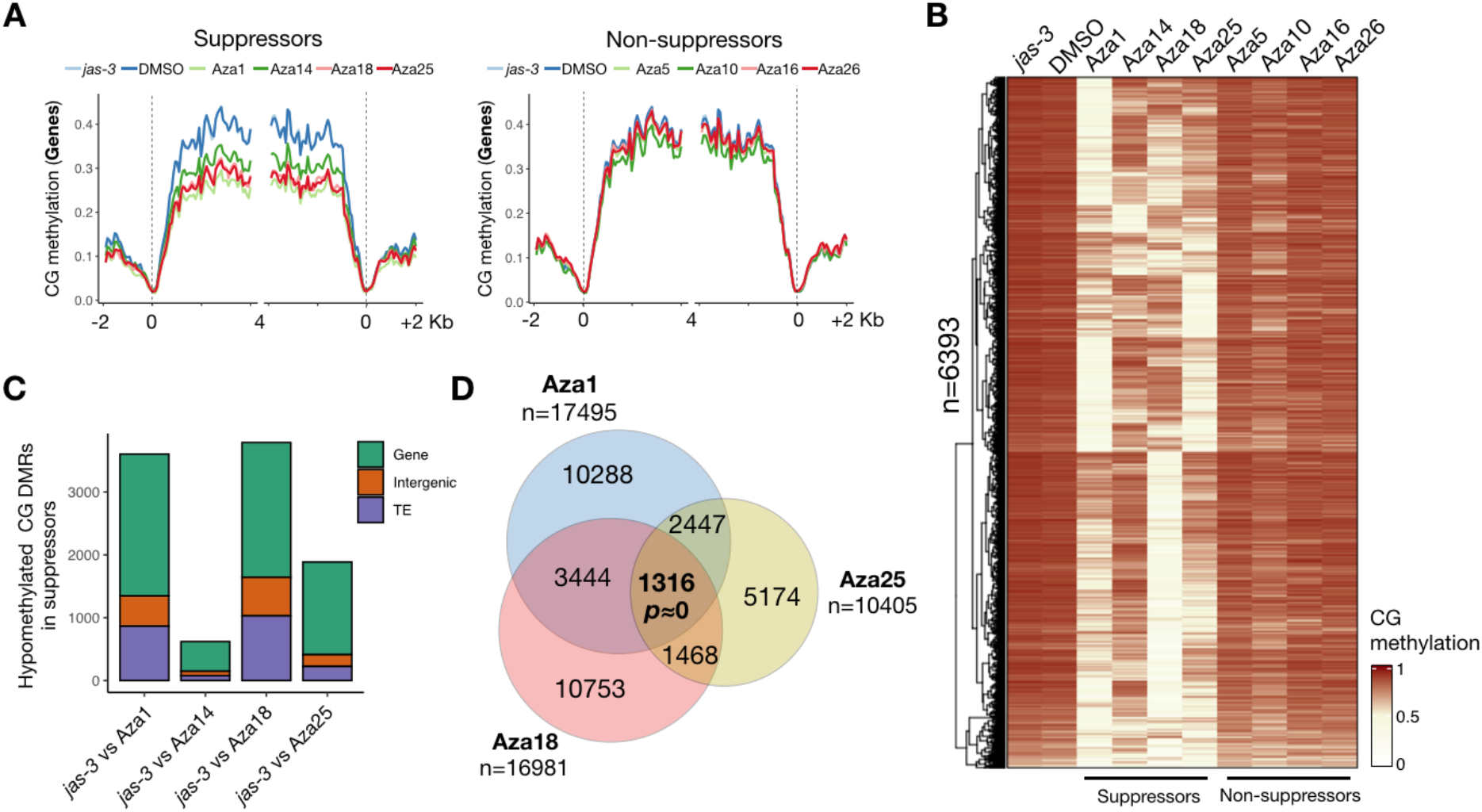
Transgenerational inheritance of CG hypomethylation occurs specifically in suppressor lines. **(A)** Metaplots show CG methylation levels at protein-coding genes annotated according to the TAIR10 reference genome, and aligned at the 5’ and 3’ ends (dashed lines). Average CG, methylation was calculated for 100-bp intervals and plotted for untreated *jas-3* and DMSO controls, suppressor and non-suppressor lines, showing that loss of CG methylation occurs specifically in the suppressor lines. **(B)** Heatmap representation of CG methylation levels at hypomethylated CG DMRs detected in the suppressor lines (Aza1, Aza14, Aza18 and Aza25) as compared to the untreated control *jas-3*. Average CG methylation mapping to these DMRs is presented as heatmap for untreated *jas-3* (2 replicates) and DMSO controls (2 replicates), suppressors and non-suppressors. **(C)** Hypomethylated CG DMRs detected in each suppressor line were mapped to the genomic features annotated in the TAIR10 reference genome, showing that the majority of DMRs overlap with protein-coding genes and transposable elements (TEs). **(D)** Venn diagram shows a significant overlap between differentially methylated 100bp bins detected in the three strongest suppressor lines (Aza1, Aza18 and Aza25). The statistical significance of the observed regions was calculated using the R package SuperExactTest (Wang et al., 2015).

In DNA methylation mutants, such as *met1* and *ddm1*, ectopic CHG methylation is often observed in gene bodies (Stroud et al., 2013), and may lead to transcriptional gene silencing. Similarly, ectopic CHG methylation was observed at paternally expressed genes down-regulated in the endosperm of viable triploid seeds after pollination with diploid *met1* pollen (Schatlowski et al., 2014). It was hypothesized that ectopic CHG methylation at the paternal *met1* genome leads to reduced expression of PEGs during endosperm development, thus contributing to viability of seeds with paternal excess (Schatlowski et al., 2014). Although this effect was not previously reported in plants treated with 5-Azacytidine (Griffin et al., 2016), we detected 95 hypermethylated CHG DMRs in the progeny of the strongest suppressor line Aza1 (Supplemental Figure 6). This indicates that newly formed epialleles with ectopic CHG methylation are induced by 5-Azacytidine treatment, and are stably inherited to the next generation. However, this effect was not consistently observed among the four suppressor lines tested (Supplemental Figure 6), suggesting that ectopic CHG is also not responsible for the suppressive effect.

### The triploid block response and DNA methylation levels were partially restored two generations after 5-Azacytidine treatment

Epigenetic variation induced by 5-Azacytidine is known to be a transient effect (Pecinka and Liu, 2014), and only a few epialleles have been detected in subsequent generations after treatment (Akimoto et al., 2007). However, we found approximately 3000 hypomethylated CG DMRs in the eM1 generation of independent suppressor lines (Figure 2C, Supplemental Data Set 2), which confirms that hypomethylation induced by 5-Azacytidine is stable for at least one generation after treatment. In order to investigate further the transgenerational stability of the suppressive effect, two diploid eM1 siblings from the strongest suppressor Aza1 were selected and selfed (Aza1-1 and Aza1-2), and six diploid eM2 plants were analyzed individually for each line (Supplemental Figure 1). Seed abortion and germination analysis showed clear signs of recovery of the triploid block response in both lines, although only Aza1-2 was significantly higher compared to the levels detected in the eM1 generation (Figure 3A). We also performed WGBS analysis of bulked seedlings from each eM2 line, and confirmed that CG methylation was fully restored at many loci (Figure 3B), while ectopic CHG methylation was no longer detected (Supplemental Figure 6). This strongly suggests that a progressive recovery of CG methylation contributes to a stronger triploid block response in eM2 plants. However, additional analyses will be required to determine if DNA methylation and triploid block response are fully restored to *jas-3* levels in subsequent generations.

**Figure 3.**
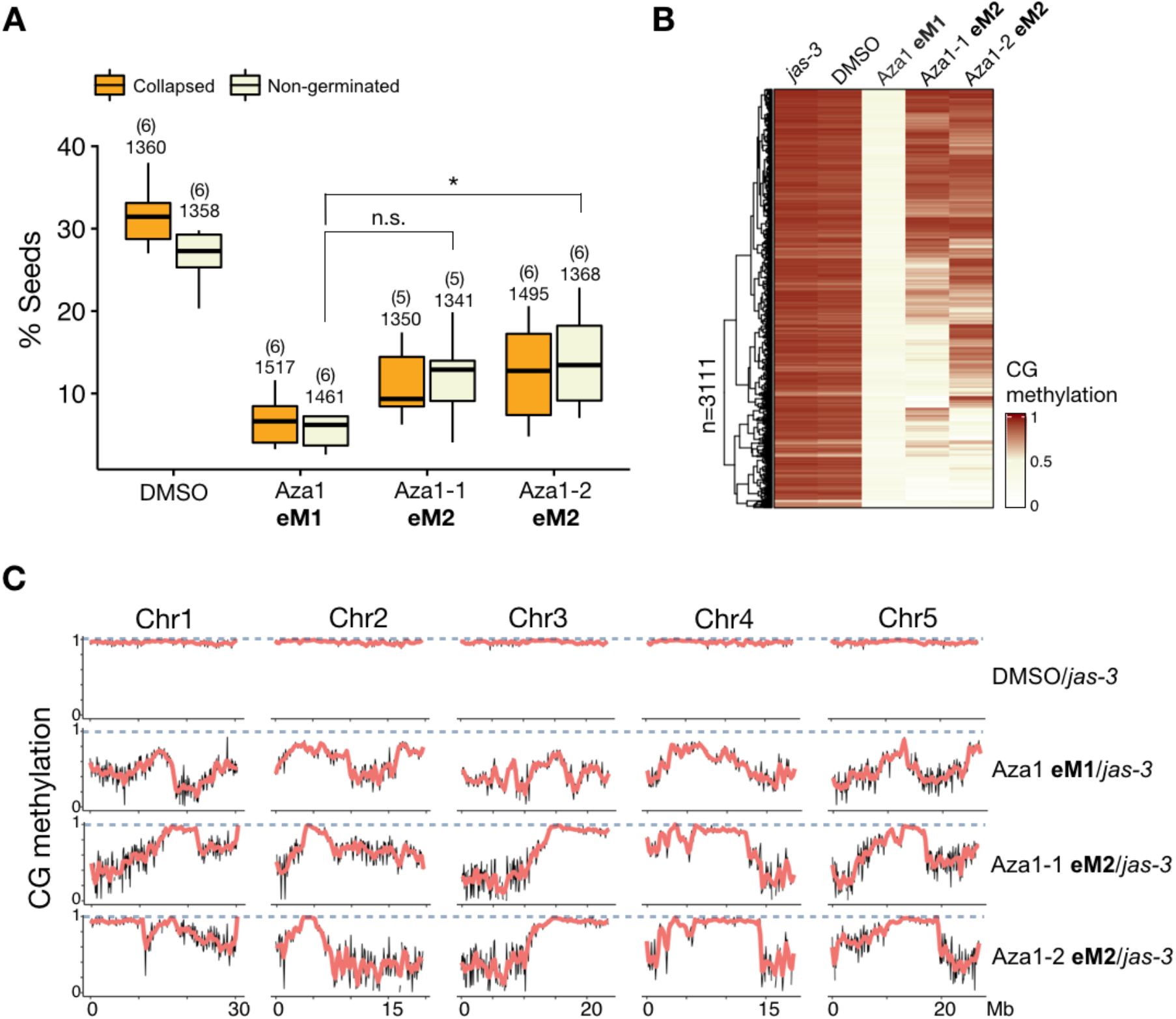
The triploid block response and CG methylation are partially restored in the second generation after treatment with 5-Azacytidine. **(A)** Seed abortion and germination assays were performed for two consecutive generations after treatment with 5-Azacytidine (eM1 and eM2), showing that the triploid block response is only partially restored in two independent lines in the second generation after epimutagenesis (eM2). Numbers on top of each box indicate the number of siblings (top) and total number of seeds (bottom) counted. Statistical significance was calculated by ANOVA with a post hoc Dunnett test, using Aza1 as the reference group (n.s. is not significant, and * is *p* ≤ 0.05). **(B)** Average CG methylation mapping to hypomethylated CG DMRs detected between the suppressor line Aza1 and the untreated control *jas-3* is presented as heatmap for untreated jas-3 (2 replicates) and DMSO (2 replicates), Aza1 eM1, and two eM2 lines (Aza1-1 and Aza1-2), to show that CG methylation is also partially restored in eM2 lines. **(C)** CG methylation of untreated *jas-3*, DMSO, eM1 and eM2 lines was mapped to 100kb bins across the Arabidopsis genome. CG methylation levels in DMSO, eM1 and eM2 datasets were then divided by the levels in *jas-3*, and plotted separately for all five chromosomes to show patterns of hypomethylation. This shows that for most of the genome, the distribution of CG hypomethylation is identical between the two eM2 lines, although there are particular loci where CG methylation is segregating.

Interestingly, some DMRs segregated in the eM2 generation, including a large region in chromosome 1 that restored CG methylation almost completely in Aza1-2, but remained hypomethylated in Aza1-1 (Figure 3C). Perhaps such differences explain the higher levels of triploid block in Aza1-2 (Figure 3A), although the majority of DMRs in all five chromosomes restored CG methylation to similar levels in both lines (Figure 3B and 3C), thus indicating that certain epigenetic states were already fixed. Further analyses with methylomes of additional lines and in different generations will be required for a robust evaluation of epiallele segregation. However, since the triploid block response remained relatively low in eM2 lines as compared to DMSO controls (Figure 3A), we can conclude that DMRs that fully restored CG methylation in eM2 are likely not major contributors for the suppressive effect observed in the eM1 generation.

### Paternally expressed genes are transiently down-regulated in suppressor lines

Previous studies in Arabidopsis and crops have shown that the triploid block leads to striking changes in the gene expression program of the developing triploid seed (Schatlowski et al., 2014; Stoute et al., 2012). Most notably, expression of many PEGs is up-regulated in abortive triploid seeds, and restored to wild-type diploid levels when the triploid block is genetically or epigenetically suppressed (Schatlowski et al., 2014; Kradolfer et al., 2013; Batista et al., 2019; Satyaki and Gehring, 2019; Martinez et al., 2018; Wolff et al., 2015). Therefore, we performed transcriptome analysis by mRNA sequencing of developing siliques collected 6 to 9 days post anthesis, as previously described (Mizzotti et al., 2018). Comparisons between wild-type Col-0 (WT) and *jas-3* mutant siliques allowed to detect 668 down-regulated and 1804 up-regulated genes in *jas-3* (Figure 4A, Supplemental Data Set 3). Similar analysis between WT and DMSO control plants showed a strong overlap between genes up- and down-regulated in untreated *jas-3* and DMSO control plants (Supplemental Figure 7 and Supplemental Data Set 3). These observations, together with a strong correlation between *jas-3* and DMSO replicates (Supplemental Figure 8), show that DMSO treatment does not induce major changes in the transcriptome of developing siliques.

**Figure 4.**
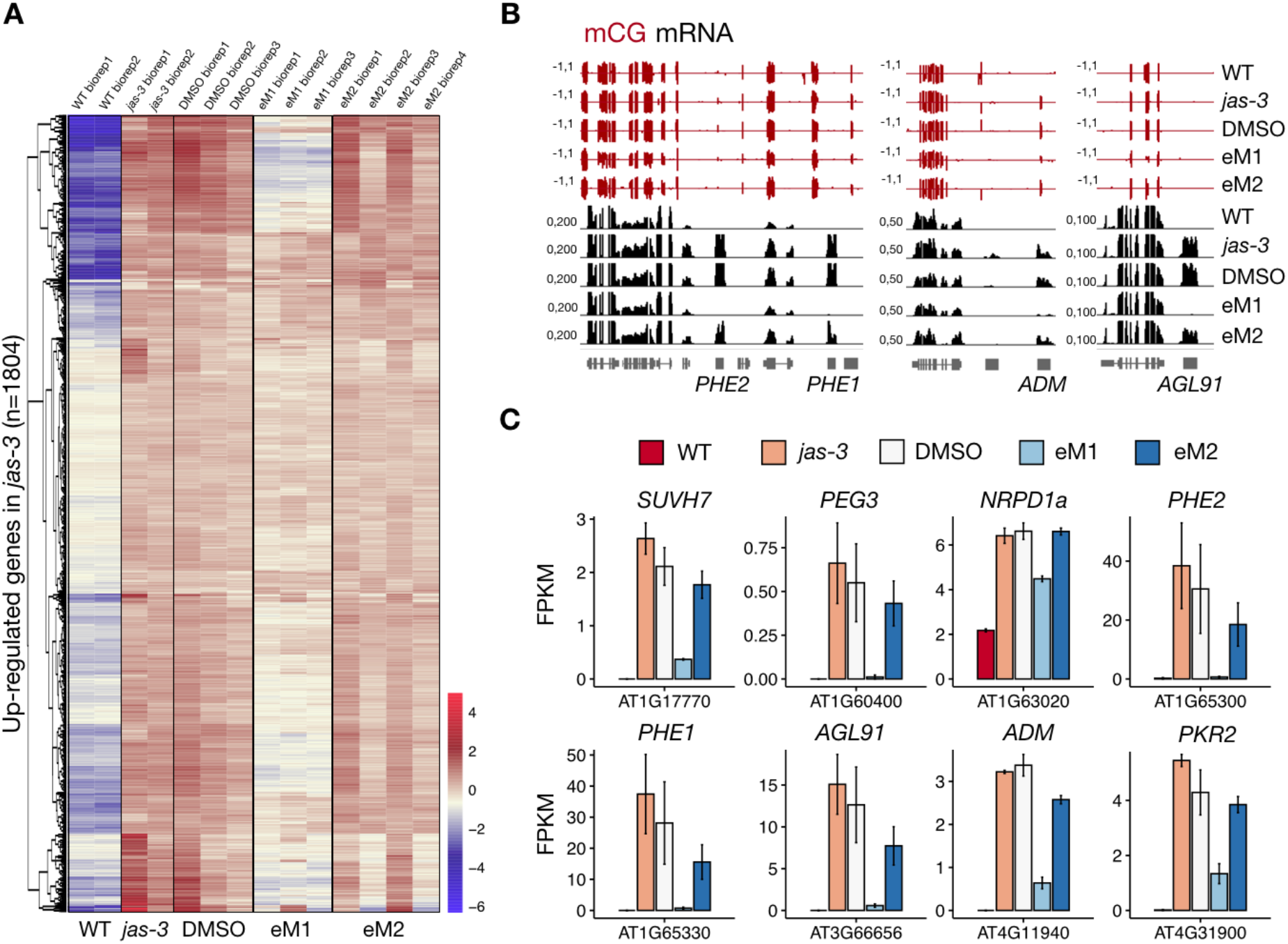
Paternally expressed genes are down-regulated in suppressor lines. **(A)** Differential expression analysis was performed between wild-type Col-0 (WT) and *jas-3* mutant siliques, and 1804 genes were found up-regulated in *jas-3* (fold change ≥ 2, adjusted *p*-value < 0.01). Transformed raw counts for these genes are presented as an heatmap for WT, *jas-3* and DMSO, Aza1 eM1 and eM2 biological replicates (see Methods), to show that a proportion of genes up-regulated in untreated *jas-3* and DMSO controls are transiently down-regulated in the first generation after treatment with 5-Azacytidine. **(B)** Genome browser tracks display CG methylation (pooled seedlings) and mRNA levels (siliques) in *jas-3*, DMSO, Aza1 eM1 and Aza1 eM2 plants. **(C)** Raw counts were normalized as fragments per kilobase per million (FPKM) and plotted as barplots with mean values and standard error.

We then compared suppressor Aza1 plants from eM1 generation with DMSO controls, and found 470 genes down-regulated in eM1, while only 48 were up-regulated (fold change ≥ 2, *p* < 0.01) (Supplemental Data Set 1). As expected, the vast majority of genes down-regulated in Aza1 eM1 plants (93%) overlapped with genes up-regulated in *jas-3* (vs WT) (Supplemental Figure 7). We also performed transcriptome profiling in siliques of four different siblings in the eM2 generation, as CG methylation and the triploid block response were partially restored in these plants (Figure 3). Indeed, we found 368 genes significantly up-regulated in eM2 siliques as compared to eM1, while only 28 genes were found down-regulated in eM2 (fold change ≥ 2, *p* < 0.01). Strikingly, most of the 368 genes up-regulated in eM2 (90%) overlapped with the genes down-regulated in eM1 plants (vs DMSO) (Supplemental Figure 7), and are thus strong candidates to explain triploid block suppression in plants treated with 5-Azacytidine. Indeed, well-known PEGs involved in the triploid block response were found among the down-regulated genes in the eM1 generation, such as *PHE1/PHE2* and *ADM* (Figures 4B and 4C) (Batista et al., 2019; Kradolfer et al., 2013), for which expression is partially restored in eM2 (Figures 4B and 4C).

### Chemically-induced hypomethylation allows bypassing interploidy and interspecific hybridization barriers

Endosperm-based hybridization barriers are frequently observed during interploidy and interspecific crosses in a variety of plant species (Lafon-Placette and Köhler, 2016), and we asked if genome-wide epimutagenesis induced by 5-Azacytidine could suppress hybridization barriers in crosses between plants of different ploidy or between different species. To test this hypothesis, we first treated tetraploid Arabidopsis plants in Col-0 background with 100μM 5-Azacytidine during seed germination. In agreement with the results in *jas-3* mutants, we obtained 40% of viable triploid seeds when using pollen from 5-Azacytidine-treated tetraploid plants to pollinate non-treated diploid Col-0 plants (Figure 5A), while only 10% of triploid seeds were viable in the control cross using pollen from DMSO-treated plants. These results show that triploid block suppression after treatment with 5-Azacytidine is not dependent on the method used to produce diploid pollen.

**Figure 5.**
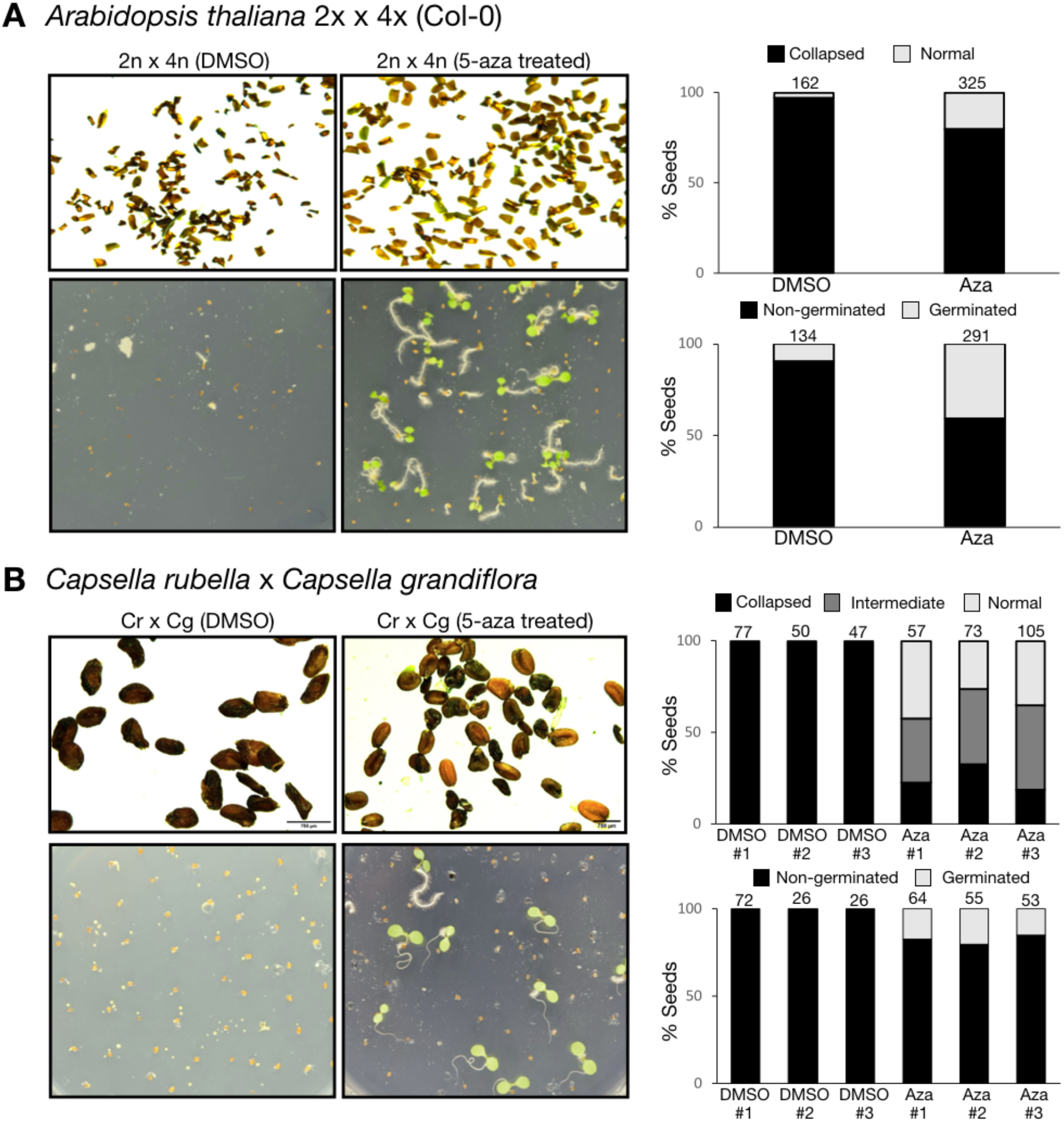
Chemically-induced epimutagenesis allows bypassing interploidy and interspecific hybridization barriers. **(A)** Pollen from tetraploid Arabidopsis plants in Col-0 background was used to pollinate diploid Col-0 plants. Triploid seeds resulting from this cross abort at high frequencies when the paternal parent is treated with DMSO, while 40% of viable triploid seeds were detected when pollen was derived from plants that have been treated with 5-Azacytidine. **(B)** In interspecific crosses between diploid *Capsella rubella* (*Cr*) and *Capsella grandiflora* (*Cg*), hybrid seeds abort, resembling interploidy hybridizations with paternal excess. When *C. grandiflora* plants are treated with 5-Azacytidine, approximately 20% of hybrid seeds germinated.

We then performed interspecific hybridizations between *Capsella rubella* and *Capsella grandiflora*, as crosses between these species resemble the triploid block when *C. grandiflora* is used as the male parent (Rebernig et al., 2015). *C. grandiflora* plants were treated with 100μM 5-Azacytidine, and pollen from treated plants was used to pollinate non-treated *C. rubella* plants. Strikingly, approximately 30% of hybrid seeds were plump and appeared normal, and about 40% of seeds were less severely collapsed, as compared to the control DMSO crosses that resulted in 100% completely collapsed seeds (Figure 5B). When these hybrid seeds were tested in germination assays, we found that about 15-20% of hybrid seeds could germinate in the crosses where the male parent was treated with 5-Azacytidine (Figure 5B), while none of the seeds germinated in the control cross. Collectively, our data shows that DNA hypomethylation induced by 5-Azacytidine treatment can suppress hybrid seed lethality in both interploidy and interspecific crosses.

## Discussion

The triploid block is a classic example of dosage regulation, and our recent studies have demonstrated how plants use epigenetic mechanisms to sense and control parental genome dosage in crosses between parents with differing ploidy (Martinez et al., 2018; Borges et al., 2018; Schatlowski et al., 2014; Jiang et al., 2017; Wang et al., 2018; Dziasek et al., 2021; Florez-Rueda et al., 2021; Satyaki and Gehring, 2019). Differential DNA methylation between parental genomes is essential for seed development and genomic imprinting (Adams et al., 2000; Xiao et al., 2006; Choi et al., 2002), and DNA methylation of the paternal genome is required to trigger the triploid block in Arabidopsis (Schatlowski et al., 2014; Wang et al., 2021; Satyaki and Gehring, 2019). However, and despite the recent advances, the mechanistic aspects of this complex process remain largely unclear.

In this study, we demonstrate that the triploid block is bypassed in epimutagenized plants treated with the cytosine analog 5-Azacytidine, which induces DNA hypomethylation. We found that the suppression level was highly correlated with a genome-wide and transgenerational loss of DNA methylation that occurred mostly in the CG context (Figure 2 and Supplemental Figure 4). Interestingly, independent suppressor lines showed strong loss of CG methylation at different loci, while non-suppressors had almost normal levels of CG methylation (Figure 4B). The reason for this variability among treated plants remains to be explored, but one possibility is that only a fraction of plants treated with 5-Azacytidine is able to transmit DNA hypomethylation to their germline, and from there to the next generation.

We were able to validate previous results showing that DNA hypomethylation of the paternal genome is what allows viability of triploid seeds (Schatlowski et al., 2014; Wang et al., 2021; Satyaki and Gehring, 2019). In previous studies, ectopic CHG methylation at PEGs was observed in the endosperm carrying a hypomethylated paternal genome derived from diploid *met1* pollen, and was associated with triploid block suppression (Schatlowski et al., 2014). However, only one of the suppressor lines in our study showed a small number of loci with ectopic CHG methylation (Supplemental Figure 6), thus suggesting that this mechanism is unlikely responsible for PEG suppression in 5-Azacytidine-treated plants. Nevertheless, DNA methylation analyses in the developing endosperm of different suppressor lines will be required to finally evaluate this hypothesis.

Notably, we found a significant number of DMRs overlapping between the three strongest suppressors lines Aza1, Aza18 and Aza25 (Figure 2D). It is tempting to speculate that a particular epiallele (or epialleles) involved in the triploid block is within this list of 253 DMRs (Supplemental Data Set 2), although there is no clear overlap with genes up-regulated in *jas-3,* or down-regulated in eM1 (Supplemental Data Set 3). Alternatively, triploid block suppression may simply require that a certain amount of the genome remains hypomethylated, independent of location. Comparisons between plants in eM1 and eM2 generations support this idea, as DNA methylation was restored at only a fraction of hypomethylated CG DMRs detected in eM1 (Figure 3B), but the majority of genes down-regulated in eM1 were significantly up-regulated in eM2 (Supplemental Figure 7). This way, genome-wide CG methylation levels of the paternal genome could somehow function as a “ploidy sensor” in the developing endosperm, by attracting or repulsing epigenetic modulators of genomic imprinting. The most obvious candidates for this interplay are the paternal DNA methylation and maternal PRC2 pathways that are often mutually exclusive (Deleris et al., 2012; Rougée et al., 2020), and have been independently implicated in the triploid block (Erilova et al., 2009; Martinez et al., 2018; Satyaki and Gehring, 2019; Wang et al., 2021).

Our work furthermore shows that chemically-induced epimutagenesis allows bypassing interspecific hybridization barriers in crosses between the Capsella species *C. rubella* and *C. grandiflora* (Figure 5B). Interestingly, CHG and CHH methylation was shown to decrease in Capsella hybrid endosperm, while CG methylation increased (Dziasek et al., 2021). However, it remains to be explored whether the increase in CG methylation is what causes hybrid seed defects in interspecific crosses, rather than the decrease in CHG and CHH methylation.

In conclusion, our study demonstrates that 5-Azacytidine can be successfully used as a tool to facilitate the generation of triploid seeds and interspecific F1 hybrids in different plant systems. We believe this method could be applicable to a wide range of species, including crops with high agronomical interest, thus providing a convenient and cheap strategy to facilitate modern plant breeding.

## Methods

### Plant growth and chemically-induced epimutagenesis

The mutant allele *jas-3* (SAIL_813_H03, Col-0 background) was used in this study. Diploid seeds from *jas-3* mutants were surface sterilized with 50% bleach for five minutes, rinsed with sterile deionized water, and sowed on agar plates containing 0.5X Murashige and Skoog (MS) medium, 1% sucrose, pH = 5.7 and different concentrations (25, 50 and 100μg/mL) of 5-Azacytidine (Sigma), under long day conditions (16h photoperiod). DMSO solvent was used as control treatment. Seedlings were transferred to soil after two weeks, and maintained in greenhouse long-day conditions to complete the life cycle.

### Triploid block quantification

The triploid block was quantified by counting the number of aborted seeds in 5 siliques after selfing. The same set of seeds was then surface-sterilized using 50% bleach and ethanol, rinsed once with ethanol 96%, and air-dried. Seeds were then sown on agar plates containing 0.5X MS medium, 1% sucrose, pH = 5.7, then stratified for 2 days at 4 °C, and transferred to growth chambers under long-days conditions (16h day, 8h dark). Germination rate was initially quantified on 4 to 5-day-old seedlings, then adjusted after 7 days if necessary to account for germination delays.

### Pollen ploidy analysis by flow cytometry

Pollen ploidy in *jas-3* mutants was analyzed by collecting open flowers from individual plants into eppendorf tubes, vortexing in 2mL of 100mM sodium phosphate buffer (pH 7) for 3min, and filtering through a 50μm nylon mesh. Pollen population is characterized by an elevated high angle scatter (SSC) and autofluorescence, which allows discrimination of haploid (1n) and diploid (2n) pollen, as previously described (Storme and Geelen, 2011; Erilova et al., 2009). These two populations were gated and quantified (Supplemental Figure 2). For ploidy analysis of nuclei, leaf tissue was chopped in 2mL of Galbraith buffer (45 mM MgCl2, 20 mM MOPS, 30 mM sodium citrate, 1 % (v/v) Triton X-100. Adjust pH to 7.0) using a razor blade, filtered through a 50μm mesh, stained with SYBR Green dye (Lonza), and analyzed on a CyFlow Space cytometer (Sysmex).

### Whole-genome bisulfite sequencing and DNA methylation analysis

Bulked seed from each Aza line, two *jas-3* and two DMSO plants were germinated on MS plates, genomic DNA was isolated from ten pooled seedlings using the Quick DNA purification kit (Zymo), and library preparation and sequencing was performed by BGI Genomics (Hong Kong). Briefly, genomic DNA was fragmented by sonication, end-repaired and ligated to methylated adaptors. After bisulfite treatment, bisulfite-treated fragments were PCR amplified and sequenced as paired-end 100bp reads (PE100) with DNBSEQ technology (BGI). Pre-processed and high-quality reads were mapped to the TAIR10 genome using bismark with default settings for paired-end libraries (Krueger and Andrews, 2011), and all figures and downstream analysis were performed using R. DMRs in CG context were defined as 100-bp bins containing at least 4 differentially methylated CGs, and with an absolute methylation difference of 0.4. DMRs localizing within 200 bp of each other were merged. The statistical significance of the observed overlaps between differentially methylated 100bp bins was calculated using the R package SuperExactTest (Wang et al., 2015). A summary of all bisulfite sequencing data generated in this study is presented in Supplemental Data Set 1, and is accessible through the NCBI’s Gene Expression Omnibus (GSE179702).

### RNA sequencing and analysis

Total RNA was extracted from three siliques 6-9 days after anthesis, as previously described (Mizzotti et al., 2018), and using the RNeasy Plant Mini Kit (Qiagen) following manufacturer’s recommendations for seed tissues (RLC buffer). Sequencing of messenger RNA was performed by BGI Genomics (Hong Kong) using DNBSEQ technology. High quality raw reads were aligned to the TAIR10 genome using STAR (Dobin et al., 2013). Reads were counted and normalized using the R package DESeq2 (Love et al., 2014). We considered differentially expressed genes those displaying a log2 fold-change ≥ 2, and with an adjusted *p*-value < 0.01. Graphical outputs were produced using the R packages ggplot2, pheatmap and ComplexHeatmap. The statistical significance of the observed overlaps between differentially expressed genes was calculated using the R package SuperExactTest (Wang et al., 2015). A summary of all RNA sequencing data generated in this study is presented in Supplemental Data Set 1, and is accessible through the NCBI’s Gene Expression Omnibus (GSE179697).

### Interploidy and interspecific hybridizations

Seeds of *Capsella rubella* (accession 48.21) and *Capsella grandiflora* (accession 23.5), as well as diploid and tetraploid Col-0 seeds were surface sterilized with 30% bleach and 70% ethanol, rinsed with distilled water and sown on agar plates containing 0.5X Murashige and Skoog (MS) medium and 1% sucrose. Seeds of *C. grandiflora* and tetraploid Col-0 were also sown on agar plates with 0.5X MS medium, 1% sucrose and 100μM 5-Azacytidine (Sigma). All plates were put in a growth chamber with long-day photoperiod (16h and 22°C light, 8h and 19°C darkness) with a light intensity of 110 μE. 7 days-old seedlings were transferred to pots filled with sterile soil and plants were grown in a growth chamber with 60% humidity and daily cycles of 16h light at 21 °C and 8 h darkness at 18°C with light intensity of 150 μE. Flower buds were manually emasculated and pollinated after 2 days. Dry seeds were stored for 30 days for “after-ripening”. They were then surface sterilized and sown on agar plates containing 0.5X MS medium and 1% sucrose. Plates were stratified for 2 days at 4°C and then moved to the growing chamber. Germination rate was scored after 7 days in the growing chamber. The experiment was done in 3 biological replicates (each replicate is offspring of different parental plants).

## Authors contributions and acknowledgments

F.B. designed the study, J.H., K.D., K.P. and F.B. performed experiments, J.H., T.W. and F.B. analyzed the data, C.K. provided materials and advised on experimental design, and F.B. wrote the manuscript with contributions from all authors.

We thank all members of our laboratories for daily discussions. This work was supported by the grant “EpiHYBRIDS” from the French National Agency of Research (ANR-19-CE12-0008) and by an installation grant provided by the BAP department of INRAE to F.B. This work was also partially supported by the EUR SPS-GSR (ANR-17-EUR-0007). The work of K.D. and C.K. was supported a grant from the Knut and Alice Wallenberg Foundation to C.K. (grant #2018-0206).

## Supplemental Data

**Supplemental Figure 1.** Quantification of diploid pollen in *jas-3* plants. (Supports Figure 1)

**(A)** The proportion of haploid (1n) and diploid (2n) pollen produced in *jas-3* mutants was quantified by flow cytometry, as pollen populations are characterized by an elevated high angle scatter (SSC) and autofluorescence (FL1).

**(B)** Boxplot was used to show the distribution of diploid (2n) pollen among individual DMSO and Aza1 plants. Number on top of the boxes represent the number of plants used. A Wilcoxon test was used to compare the mean values between the suppressor Aza1 and the DMSO control. n.s. is not significant (*p*=0.937).

**Supplemental Figure 2.** Ploidy analysis by flow cytometry. (Supports Figure 1)

Leaf tissue was chopped in Galbraith buffer with a razor blade, stained with SYBR Green dye (Lonza), and analyzed on a CyFlow Space cytometer (Sysmex). Nuclei of young diploid leaves in Arabidopsis is characterized by two prominent 2C and 4C peaks. Triploid individuals show a proportional increase in signal intensity.

**Supplemental Figure 3.** Suppression of the triploid block in plants treated with 5-Azacytidine is a paternal effect. (Supports Figure 1)

**(A)** A parental effect in triploid block suppression after epimutagenesis was investigated by performing reciprocal crosses between the strong suppressor Aza1 and DMSO control plants. Numbers on top of each bars represent the number of plants used (top) and total amount of seeds counted (bottom). Statistically significant difference in the percentage of collapsed seeds was calculated by ANOVA with a post hoc Dunnett test, using Aza1 as the reference group (n.s. is not significant, and *** is *p* ≤ 0.001).

**(B)** Representative seeds images are shown for the controls (selfed plants), and reciprocal crosses. Scale bars represent 1mm.

**Supplemental Figure 4.** Schematic depicting *jas-3* epimutagenesis and transgenerational analysis of the triploid block in suppressor lines. (Supports Figures 1, 2 and 4)

Diploid seeds from *jas-3* were treated with 100μg/mL of 5-Azacytdine during germination and early growth. Treated eM0 plants were then transferred to soil to recover, and allowed to self-fertilize. Selfed *jas-3* mutants produce diploid seeds that are viable, and triploid seeds that abort. Diploid plants were selected after each generation and allowed to self-fertilize, and the triploid block was quantified by counting the number of aborted seeds in individual siblings.

**Supplemental Figure 5.** CG, CHG and CHH methylation profiles at protein-coding genes and transposable elements. (Supports Figure 2)

Protein-coding genes (left panels) and transposable elements (TEs, right panels) annotated according to the TAIR10 reference genome were aligned at the 5’ and 3’ ends (dashed lines), and average CG, CHG and CHH methylation levels for 100-bp intervals were plotted for untreated *jas-3* and DMSO controls, suppressor and non-suppressor lines. Loss of DNA methylation at genes and TEs occurs mainly in the CG context and is observed only in suppressor lines.

**Supplemental Figure 6.** Ectopic CHG methylation in suppressor lines. (Supports Figure 2)

**(A)** Heatmap representation of CHG methylation levels at ectopic CHG DMRs detected in the suppressor line Aza1, as compared to the untreated control *jas-3*. Average CHG methylation mapping to these DMRs is presented as heatmap for two independent replicates of untreated *jas-3* and DMSO controls, suppressor and non-suppressor lines.

**(B)** Genome browser tracks show DNA methylation at CG and CHG context for selected loci showing ectopic CHG methylation.

**Supplemental Figure 7.** Differentially expressed genes in suppressor lines. (Supports Figure 4)

**(A)** Venn diagram shows overlap between genes up-regulated in untreated jas-3 (vs WT), and genes up-regulated in DMSO controls (vs WT).

**(B)** Venn diagram shows the overlap of genes up-regulated in *jas-3* (vs WT) with genes transiently down-regulated in eM1 (vs DMSO), and up-regulated in eM2 (vs eM1). The majority of genes down-regulated in eM1 are genes that are up-regulated in *jas-3* mutant siliques, and expression of these genes is restored in eM2.

The statistical significance of the observed overlaps between differentially expressed genes was calculated using the R package SuperExactTest (Wang et al., 2015).

**Supplemental Figure 8.** Clustering of RNA-seq datasets. (Supports Figure 4)

Transformed data was used for sample clustering to visualize sample-to-sample distances (see Methods). The heatmap of this distance matrix shows an overview over similarities and dissimilarities between all RNA-seq datasets produced in this study.

**Supplemental Data Set 1.** Summary whole-genome bisulfite and RNA-seq datasets.

Whole-genome bisulfite sequencing (WGBS). Mean genomic coverage and methylation percentages for CG, CHG and CHH context. Cytosines covered by 3 or more reads were used to calculate genome-wide methylation percentages.

RNA-seq. High quality raw reads were aligned to the TAIR10 genome using STAR (Dobin et al., 2013)

**Supplemental Data Set 2.** Lists of differentially methylated regions.

Differentially methylated regions (DMRs) in CG context were defined as 100-bp bins containing at least 4 differentially methylated CGs, and with an absolute methylation difference of 0.4. DMRs localizing within 200 bp of each other were merged.

**Supplemental Data Set 3.** Lists of differentially expressed genes.

Reads counts were normalized using the R package DESeq2 (Love et al., 2014). We considered differentially expressed genes (DEGs) those displaying a log2 fold-change ≥ 2, and with an adjusted *p*-value < 0.01.

